# Enabling Hydrogel Coating on Silicone Breast Implants with Poly(Vinyl Acetate) Primer Layer

**DOI:** 10.1101/2024.12.16.628176

**Authors:** Katrin Stanger, Dardan Bajrami, Peter Wahl, Fintan Moriarty, Emanuel Gautier, Alex Dommann, Kongchang Wei

**Affiliations:** Division of Plastic and Hand Surgery, Cantonal Hospital Winterthur, Winterthur, Switzerland; Laboratory for Biomimetic Membranes and Textiles, Empa, Swiss Federal Laboratories for Materials Science and Technology, Lerchenfeldstrasse 5, 9014 St. Gallen, Switzerland; ZHAW School of Engineering, Technikumstrasse 71, Winterthur, Switzerland; Division of Orthopaedics and Traumatology, Cantonal Hospital Winterthur, Winterthur, Switzerland; Faculty of Medicine, University of Bern, Bern, Switzerland; AO Research Institute Davos, Davos, Switzerland; Department of Orthopaedics, HFR Fribourg – Cantonal Hospital, Fribourg, Switzerland; OrthoTrauma Foundation, Fribourg, Switzerland; ARTORG Center for Biomedical Engineering Research, University of Bern, Bern, Switzerland; Laboratory for Biointerfaces, Empa, Swiss Federal Laboratories for Materials Science and Technology, Lerhen- feldstrasse 5, 9014 St. Gallen, Switzerland

## Abstract

Implant-associated infection is a major cause for breast implant re-operation. A practical method to reduce this risk is yet to be established. Hydrogel coating represents one promising approach. However, the adhesion of the hydrogel layer onto the silicone implant surface presents a significant challenge due to the intrinsic hydrophobicity of silicone surfaces. In this study, we described a surface-priming strategy involving poly(vinyl acetate) (PVAc) polymers to facilitate hydrogel adhesion to silicone implant surfaces. Miniature silicone implants with identical surface properties to clinical implants were custom-made for this study. We demonstrated that a PVAc primer layer can easily be deposited on the implant surface via a dip-coating procedure. The wettability of the implant surface was increased by this primer layer, as confirmed by contact angle measurements. The improved wettability allowed the application of a model hydrogel precursor solution (alginate) on the primed implant surface. The effectiveness of such a priming strategy in facilitating hydrogel coating was validated by testing two commercially available hydrogels on the silicone implant surface. Specifically, DAC (Defensive Antibacterial Coating) and Coseal hydrogels, representing paintable and sprayable hydrogels respectively, were successfully coated on the primed surface, as confirmed by ATR-FTIR analysis. Our surface priming strategy, which avoids surface treatments like chemical reactions and plasma irradiation that are impractical for clinical use, opens up new opportunities for exploring intraoperative hydrogel applications on silicone implants.

## Introduction

Silicone breast implants have been engineered not only for aesthetic reasons for breast augmentation, but also for reconstructive purposes after surgical breast cancer treatment. ^1, 2^ The number of implantations is increasing tremendously. In the USA alone, about 300’000 women underwent augmentation mammoplasty in 2019 for aesthetic reasons, and an additional 88’000 women underwent breast reconstruction with implants. ^3^ However, up to 41% of the implants need re-operation, either for overt septic reasons or due to capsular fibrosis, which is most likely due to low-grade infections.^4-6^ Reported incidence of infection range from less than 10% to over 60%.^4-6^ Alternatives for breast reconstruction after mastectomy include technically challenging operations, such as microsurgical autologous free flap transplantation. However, these operations bear major morbidity and complication rates.^7^ Reconstruction of the breast with silicone implants is a preferable alternative because of technical simplicity, quicker recovery time, and enhanced aesthetic outcome.^8^

Implants are most susceptible to colonization by microorganisms at implantation or within the following hours. ^9, 10^ Treatment of established implant-associated infections is expensive and cumbersome, varying from long-term antibiotic therapy to surgical debridement and implant replacement.^11^ Thus, preventing intraoperative contamination is essential.^12^ Altering the surface properties of the implant has been demonstrated to be an effective approach to reduce the risk of post-implantation complications.^13-15^ Such surface modifications include coatings with antimicrobial agents, ^16, 17^ nanomaterials,^18, 19^ adhesive interfaces,^20^ and plasma-induced chemical changes of molecular structures.^21,22^

As water-retaining soft materials made from hydrophilic polymers,^23^ hydrogels have been shown to be promising in preventing bacterial adhesion at an early stage of infection, for instance, of wounds and bone implants.^23-30^ Before degradation, they can hinder adherence of microorganisms on the surface of the implant just long enough for the immune system to eliminate any intraoperative contaminant and thus prevent biofilm formation and development of infection.^31-34^ Recent studies have demonstrated the effectiveness of hydrogel coatings in preventing biofilm development on joint replacement and fracture-fixation implants, highlighting the potential clinical utility of this approach for improving patient outcomes in breast implant surgery.^29, 35-37^ Silicone breast implants are particularly well suited for applying a dissolving coating like hydrogels, as they require no tissue integration for proper function.

The key challenge for coating silicone breast implants with hydrogels is the poor surface wettability of such hydrophobic materials. ^38^ Various strategies have been explored to improve wettability, including surface activation and oxidation to modify the surface polarity.^39^ However, such methods rely on the alteration of surface chemical structures, which could potentially raise safety concerns, as well as challenges in meeting product integrity and regulatory standards. Therefore, there is a need for easily applicable, yet effective approaches for improving silicone implant wettability. Recently, Cheng et al. utilized poly(vinyl acetate) (PVAc) as a primer layer between silicone rubber and an in-situ formed tough hydrogel to achieve hydrogel coating via photo-initiated grafting.^40^ The PVAc layer served as a bridging matrix to immobilize the tough hydrogel, formed by polymerization of synthetic monomers in the presence of initiators and crosslinkers.^40^ Due to its biocompatibility and biodegradability, PVAc has been used in various biomedical applications.^41-43^ Such a surface priming strategy offers a straight-forward approach to bond in situ formed hydrogels to silicone rubber surfaces, without altering its molecular structure. It’s effectiveness for the application of commercially available hydrogels for implantation on silicone implant surfaces is, however, yet to be verified. In this study, we explored the application of a PVAc primer layer in improving wettability of silicone breast implant surface. Successful coating of commercially available hydrogels of different types, namely a paintable physical hydrogel (DAC) and a sprayable chemical hydrogel (Coseal), on PVAc-primed implants was demonstrated, without involving additional chemicals (e.g. initiators and cross-linkers),^40^ thus providing a promising approach in the operation room to use hydrogels as coatings of silicone implants for prevention of infection.

## Materials and Methods

### Materials

Silicone implants corresponding to clinical breast augmentation implants (Motiva Implants with SmoothSilk surface, Establishment Labs Holdings Inc., Alajuela, Costa Rica) were miniaturized to a diameter of 15 mm and a height of 5 mm (Figure 1A-a), and then tested by the manufacturer following ISO-14607:2018 TS-17-026.R (Rheology Testing of Breast Implant Gels with the BTC-2000). The core is made of heated liquid silicone, which turns into a soft jellylike material with high viscoelasticity and cohesive strength after cooling. The surface is made of polymerized nanotextured silicone. Sodium alginate (Sigma Aldrich, St. Louis, MO, USA, Mw. 20 kDa) was dissolved in deionized (DI) water at room temperature (23 °C) under stirring to give a 5 wt% solution. Poly(vinyl acetate) (PVAc, Mw. 100 kDa, Sigma Aldrich) was dissolved in acetone at 40 °C under stirring to give a 2 wt% solution. Defensive Antibacterial Coating (DAC, Novagenit, Mezzolombardo, Italy) and Coseal surgical sealant (Baxter, Glattpark, Switzerland) were used as purchased (without additional drug loading). DAC is composed of hyaluronic acid (HA) and poly-D,L-lactide (PDLLA), while Coseal is composed of two different functionalized poly(ethylene glycol) (PEG) polymers, namely tetra-NHS-derivatized PEG and tetra-thiol-derivatized PEG. ^36^

**Figure 1:**
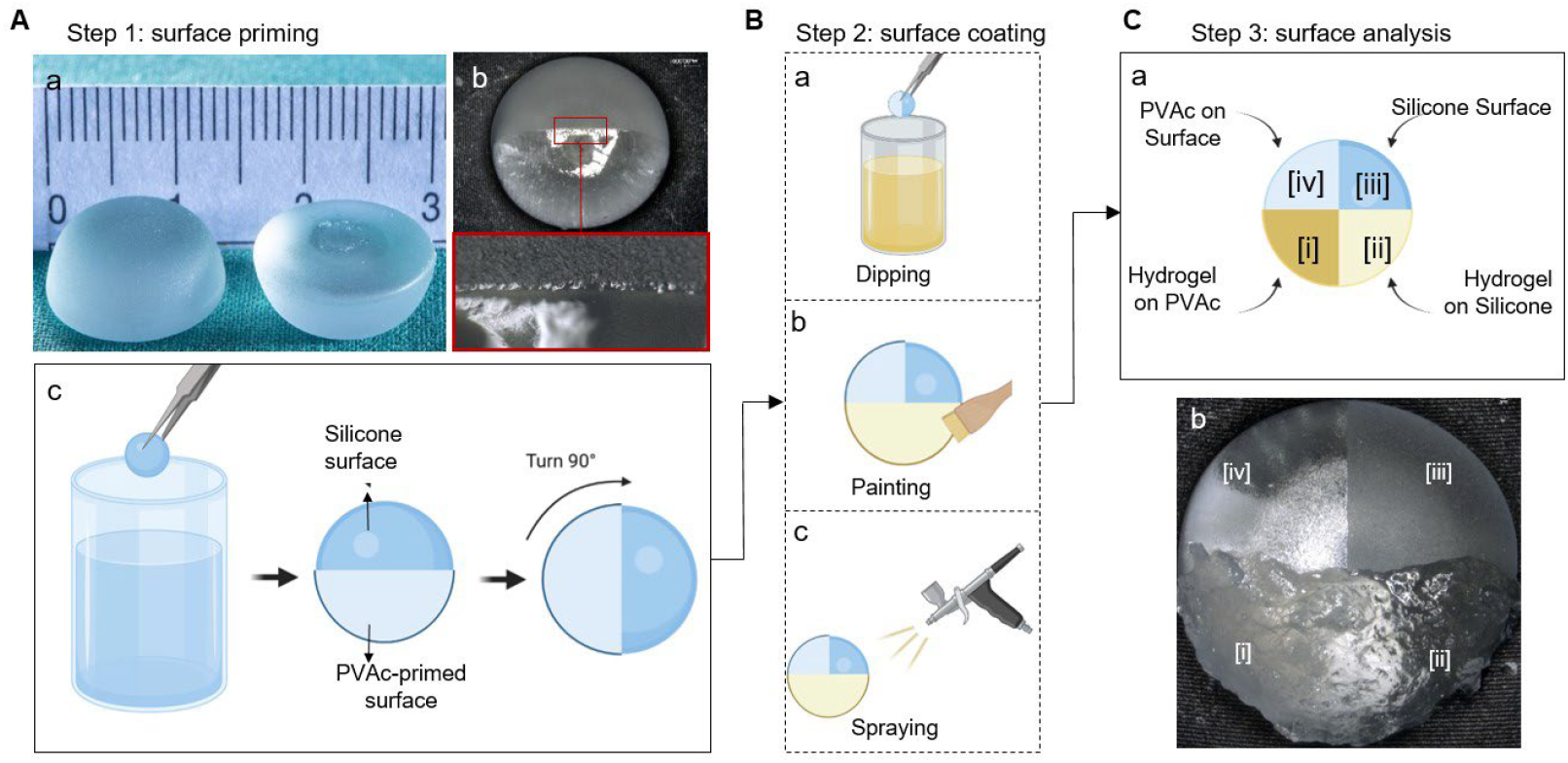
Surface priming, coating and analysis of miniaturized silicone breast implants. A-a: Custom-made silicone implants. A-b: Half of the implant primed with PVAc (upper), the boundary between the PVAc-primed and the original implant surface; A-c: PVAc coating process by dipping and rotation of the implant for the further coating process. B: Three different surface coating processes for subsequent coating of implants with alginate solution (a), DAC (b), and Coseal (c). C-a: Implant samples with four zones for the analysis; C-b: A representative microscope image of coated implant with four different zones (DAC in zone [i] and [ii]).

### Surface priming

The implants were thoroughly cleaned by using a stepwise process. They were firstly soaked in acetone and isopropanol for 10 minutes each, then cleaned by ultrasonication in deionized water for 10 minutes. After drying at room temperature in air, the implants were partially dipped into the PVAc solution for 3 seconds, followed by 15 minutes drying in air. This dipping-drying cycle was repeated 3 times, as illustrated in Figure 1A-a. The samples were subjected to surface analysis after drying in air for 1 h.

### Surface coating

Two commercially available hydrogels were tested on the PVAc primed implants. The DAC hydrogel was applied to the implants using the manufacturer’s applicator (Figure 1B-b). The Coseal hydrogel was sprayed onto the implants using a spraying system provided by the manufacturer (Figure 1B-c). The spraying was controlled by covering areas that were not to be treated. In order to analyze the coating performance on surfaces with different features, the coating process was designed as illustrated in Figure 1C.

### Surface analysis

Contact angle measurements on original implant surface and PVAc-primed implant surface were conducted by using a drop shape analyser (DSA25E, KRÜSS, Hamburg, Germany) at 23°C temperature. The implants coated with alginate solution were examined under an optical microscope (VHX-1000, Keyence, Osaka, Italy) and photographed at 10-second intervals to assess the solution’s flow behavior. The implants coated with DAC and Coseal hydrogels were washed twice in phosphate-buffered saline (PBS, 0.01 M; Sigma Aldrich). Microscopic images were taken after each washing step. One hour after coating, the individual surface zones of all samples were characterized by Attenuated Total Reflectance Fourier-Transform Infrared spectroscopy (ATR-FTIR) (Varian 640-IR, Agilent Technologies, Santa Clara, USA). The dipping method chosen provides 4 zones (i: hydrogel on PVAc; ii: hydrogel on native silicone; iii: uncoated native silicone; iv: PVAc primed silicone) on the surface of each implant (Figure 1C), allowing direct comparison of the coating.

## Results and Discussion

### PVAc primer layer alters silicone surface properties

A hydrogel directly applied on the silicone implant leads to immediate retraction (Supporting Information, Figure S1). This is due to the low wettability between silicone surface and aqueous solutions.^38^ To improve wettability, we enhanced the hydrophilic surface properties by application of a PVAc primer layer on the silicone surface. To achieve comprehensive coverage over the inherently rough implant surface, the dip-coating process was repeated three times. The PVAc primer layer showed a smooth coverage on the implant surface without disruption (Figure 1A-b). Using fewer layers resulted in inadequate coverage, which caused hydrophobic rejection of the subsequently applied hydrogel layer (Supporting Information, Figure S2).

The improved wettability of the PVAc-primed implant surface was revealed by contact angle measurement (Figure 2). The untreated silicone implant (-PVAc sample) exhibited a contact angle of 107.3 ± 12.4°, indicating a hydrophobic surface. In contrast, the PVAc-primed silicone implant surface (+PVAc sample) showed increased wettability, as evidenced by the significantly reduced contact angle of 62.3 ± 1.9°. Subsequently, it was further validated by applying an alginate solution on both the primed and non-primed implant surface (Figure 1 B-a). Alginate is a natural polysaccharide found in brown algae, commonly used in wound dressings, tissue engineering, and drug delivery applications due to its biocompatibility, low toxicity, and gelation properties.^44^ Therefore, alginate was selected herein to simulate the precursors of the commercially available hydrogels. It was observed that the alginate solution applied to the non-primed implant surfaces (-PVAc) suffered from rapid retraction, as did the hydrogels. The alginate solution was found retracted partially from the implant surface (zone [ii]) in the first 10 s after dip-coating (Figure 3A). After two minutes, the alginate solution had almost entirely disengaged from the implant, with only residual material remaining on the surface (Figure 3B-D; Supporting Information, Movie S1). On the contrary, the alginate solution applied to the PVAc-primed surface (+PVAc) retained its position without any displacement or disintegration (Figure 3D, zone [i]), thus confirming the bridging effect of the PVAc primer layer between silicone surface and hydrogel precursor solutions.

**Figure 2:**
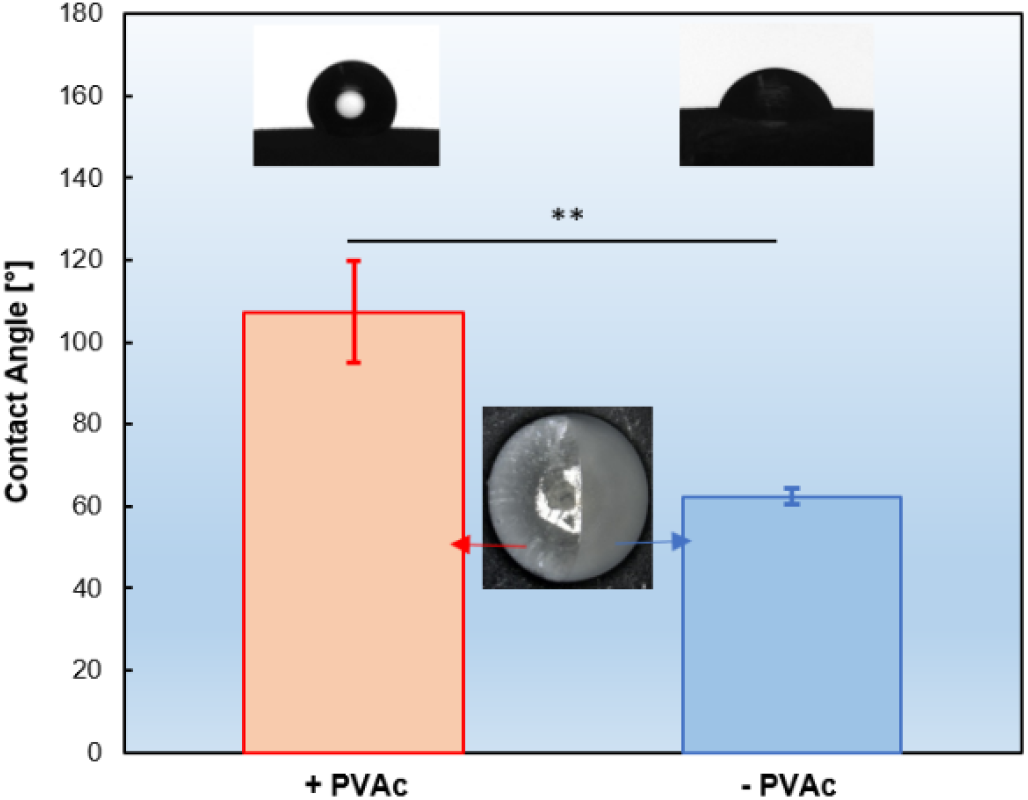
Contact angle measurements of PVAc-primed silicone implants (+PVAc) and the non-primed ones (-PVAc). Error bars represent the standard deviation of the measurements. (n = 3, ** p < 0.005)

**Figure 3:**
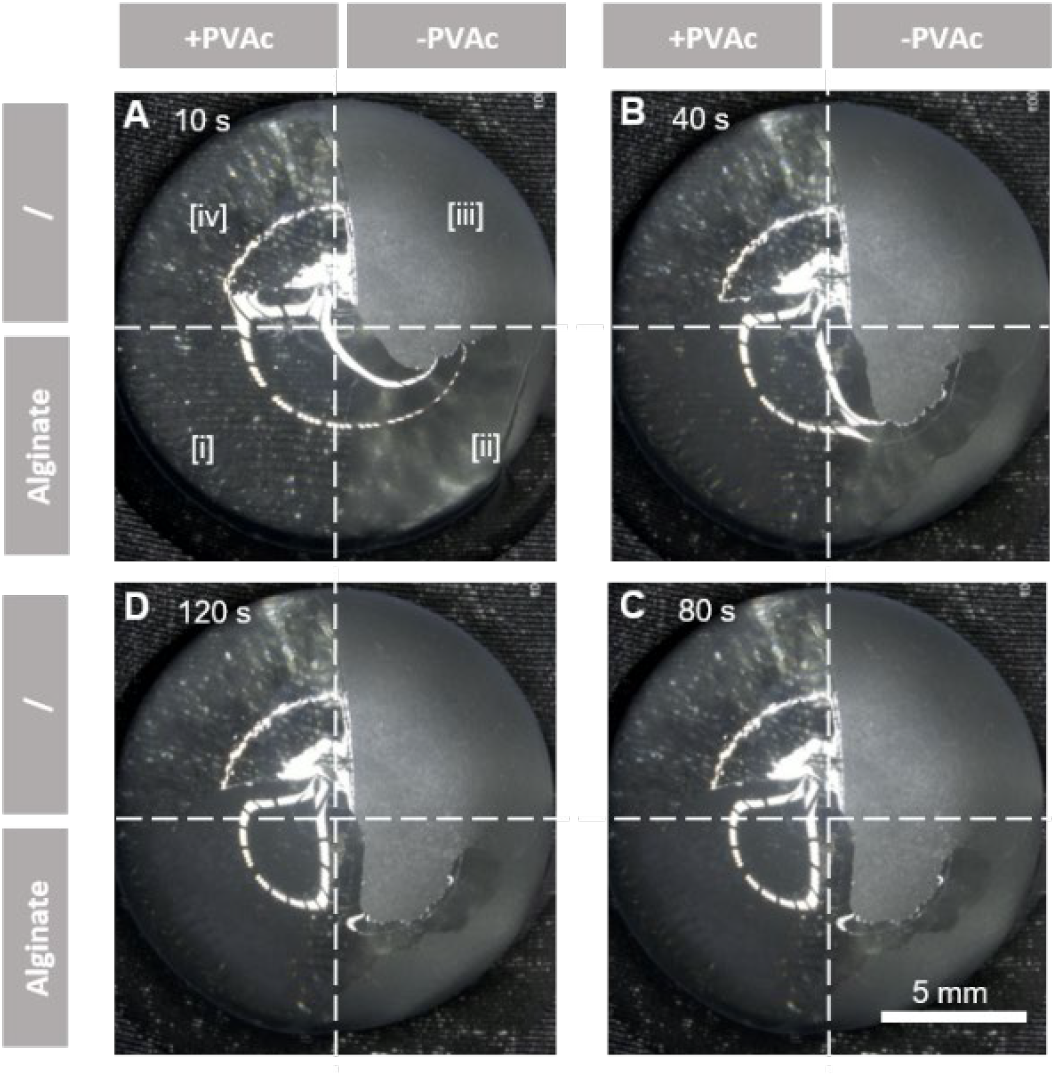
PVAc primer layer alters silicone surface properties. Silicone implants were divided into four zones, visualized by dotted white lines. The images were taken at 10s (A), 40s (B), 80s (C), and 120s (D) after dipping the implants (Zone [i] and [ii]) in alginate solution.

The change of surface property was characterized by ATR-FTIR analysis (Figure 4). On the PVAc-primed surface (zone [i]), alginate-derived signals, including the O-H stretching vibrations caused by hydroxyl groups, intermolecular hydrogen bonding at 3200-3600 cm^-1^, a sharp peak near 1600 cm^-1^ attributed to the asymmetric stretching of the carboxylate groups (COO-), and a sharp peak around 1030 cm^-1^, typically associated with C-O stretching vibrations ^45^, were observed. On the contrary, implant surface without PVAc layer (zone [ii]) lacked such signals, confirming the poor adhesion between alginate and a native silicone surface. Specifically, the peaks at 1260 cm^-1^ (Si-CH3 bending vibrations), 1010-1100 cm^-1^ (Si-O-Si stretching vibrations), and around 800 cm^-1^ (Si-C stretching vibration) revealed the bare silicone surface after unsuccessful alginate coating 26 (zone [ii]), which is in accordance with signals from the reference surface (zone [iii]). Only alginate residues were detected on such non-primed surfaces, with peaks in the 3200-3600 cm^-1^ and 1600 cm^-1^ regions (zone [ii]). It is noteworthy that when the ATR-FTIR signals from zone [ii] were compared to those from zone [iii], a difference in silicone-derived signal amplitude was obvious. This could be due to the change in surface roughness and refractive index induced by the alginate residues on the zone [ii] surface. The native implant surface with higher roughness (zone [iii]) compromised the contact between the sample and the ATR crystal, leading to weaker signals from zone [iii] than zone [ii].

**Figure 4:**
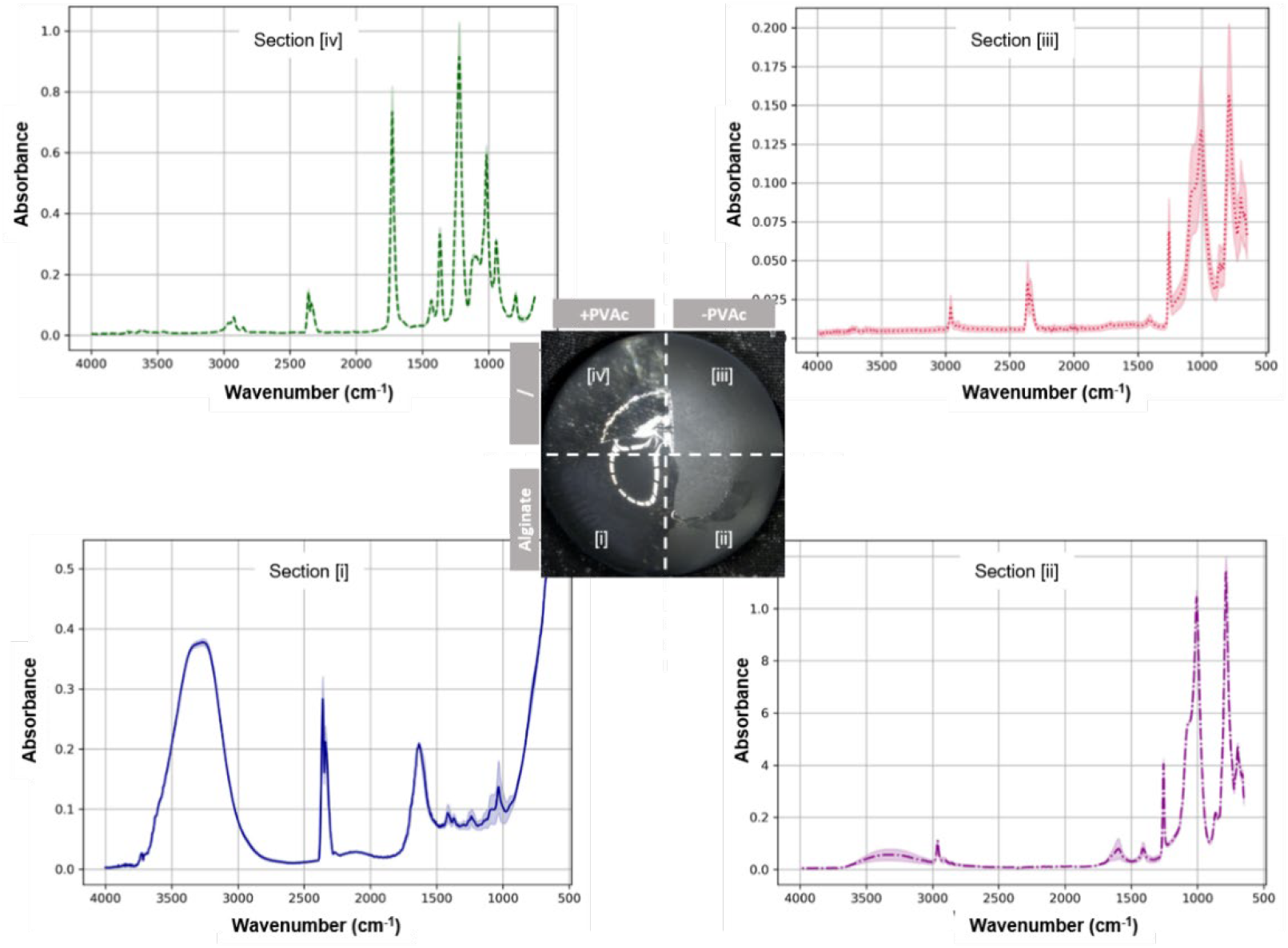
ATR-FTIR spectra of different implant surface zones measured one hour after alginate coating (n = 3).

In addition, successful priming of the implant surface with PVAc was confirmed by signals at approximately 1730 cm^-^Ü from zone [iv]) attributed to the acetate’s carbonyl, and around 1240 cm^-^Ü attributed to C-O stretch from the ester linkage, as well as peaks between 2800 and 3000 cm^-1^ for the alkyl groups (zone [iv]). According to this analysis, we confirmed that a PVAc primer layer could be applied on silicone implant surfaces without alteration of the molecular structure, offering a praticable approach to improve the wettability of the implant surface for retention of applied aqueous solutions (e.g. hydrogel precursors). Compared to other surface treatments, such as chemical activation^40^ and plasma treatment,^21, 22^ this PVAc priming approach represents a more feasible approach for intraoperative application on implants. Next, we demonstrated the successful coating of silicone breast implants with two types of commercially available medical hydrogels.

### PVAc-mediated coating of silicone implants with paintable hydrogel (DAC)

DAC is commercially available for implant surface coating, where it acts a physical barrier for infection prevention.^46^ Unlike the liquid-form alginate precursor solution, DAC is a high-viscosity physical hydrogel, which cannot be applied to the implant surface by dip-coating. As instructed by the manufacturer, DAC was applied to the implant surface with the brush provided by the manufacturer.

Before coating, the PVAc primer layer showed a homogenous and smooth surface (Figure 5A, zone [iv]). Applying a uniform layer of DAC was challenging, mainly due to the application method and the nature of the preformed high-viscosity hydrogel (zone [iii]). However, the zones [i] and [ii] can be fully coated via painting (Figure 5B). After the first wash with PBS, DAC coatings on PVAc-primed surfaces remained intact (zone [i]), although some superficial parts were flushed away (Figure 5C). On the contrary, on the untreated surfaces (zone [ii]), DAC was substantially disengaged from the implant. After the second wash, the remaining DAC hydrogel in zone [ii] was further displaced (Figure 5D). On the contrary, the hydrogel on the PVAc-primed surface (zone [i]) was still intact. This demonstrated the improved adhesion between silicone implant surface and the DAC hydrogel provided by the PVAc primer layer, thus demonstrating the stable coating of silicone implants with pre-formed DAC hydrogels. It is noteworthy that DAC residues were still observable on the interface between zones [i] and [ii], which might be stabilized by lateral forces from zone [i] to zone [ii].

**Figure 5:**
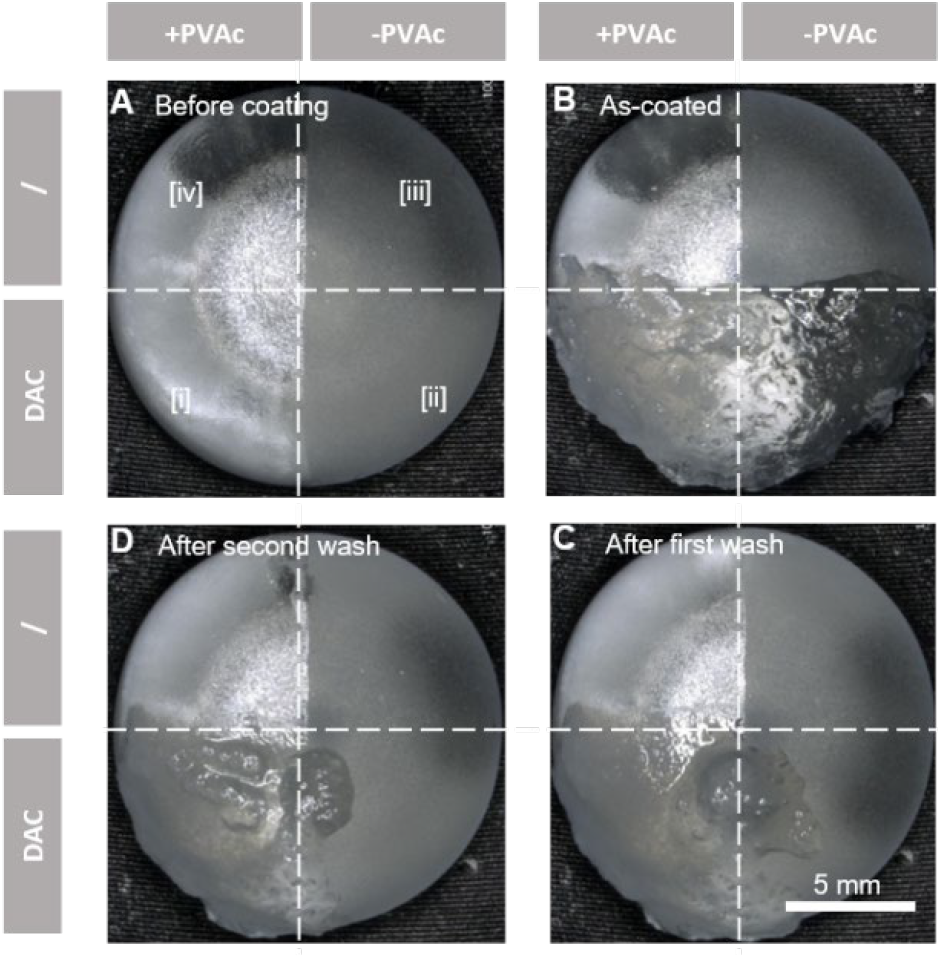
PVAc primer layer mediated coating of silicone implants with paintable DAC hydrogel. Silicone implants were divided into four zones, visualized by dotted white lines. The images were taken before coating with DAC (A), immediately after coating on zone

The ATR-FTIR difference between zone [iii] and [iv] first confirmed the successful PVAc priming. Secondly, the stable DAC coating was validated by comparing zone [i] to zone [ii]. Same peaks as those from the silicone reference surface (zone [iii]) were observed from zone [ii], indicating an absence of the DAC hydrogel (Figure 6). On the other hand, on the PVAc-primed surface, the presence of hyaluronic acid (HA) and poly-D,L-lactide (PDLLA) was confirmed (Figure 6, zone [i]).

**Figure 6:**
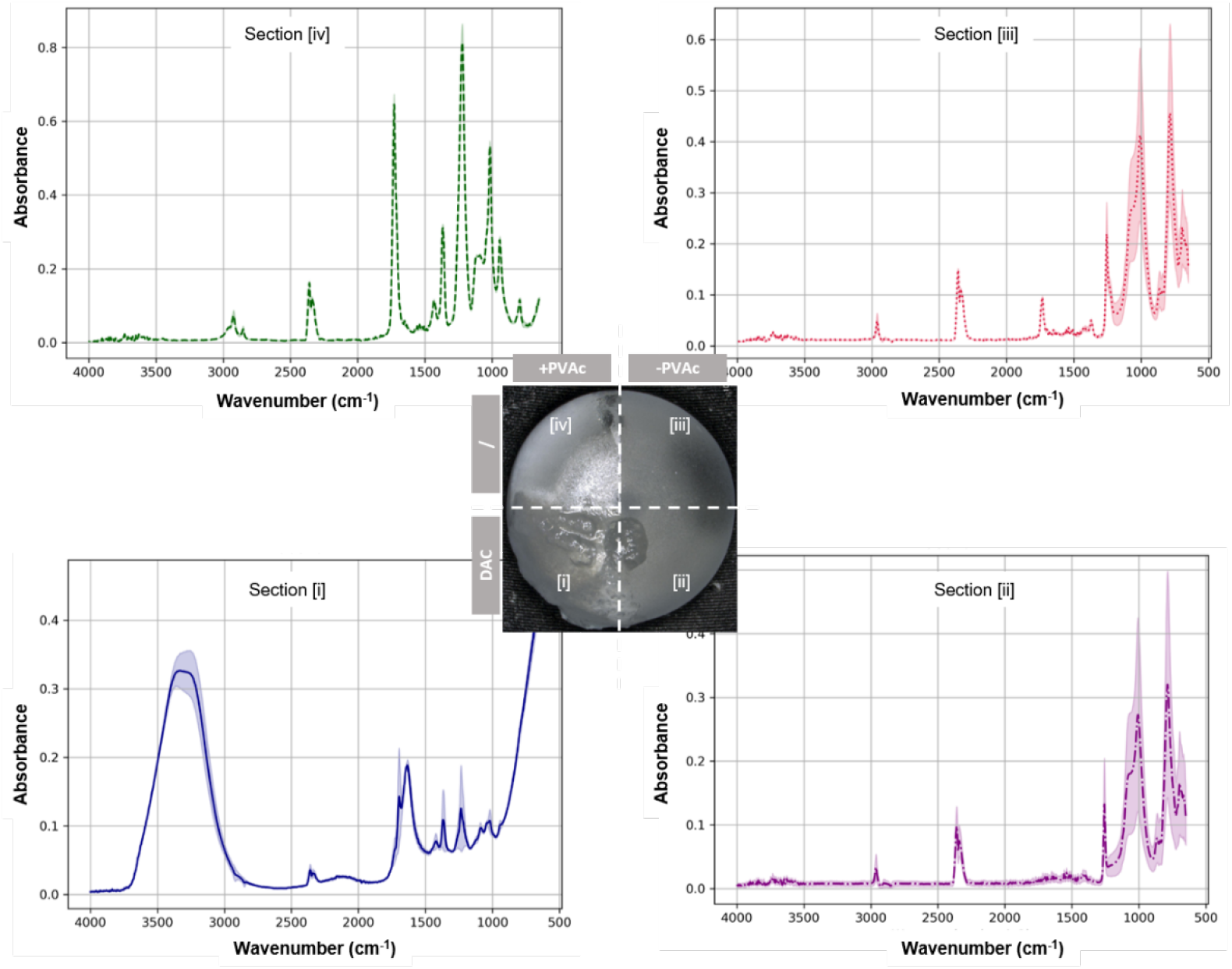
ATR-FTIR spectra of different implant surface zones after coating zone [i] and [ii] with DAC hydrogels (n = 3).

More specifically, for HA, broad bands around 3300 cm^-1^ were observed due to O-H stretching from hydroxyl groups. Additionally, a peak near 1620 cm^-1^ due to C-O stretching and another around 1400 cm^-1^ corresponding to the carboxylate ion (COO-) asymmetric stretching, as well as additional bands in the range of 1000-1100 cm^-1^ due to C-O-C stretching vibrations of the sugar backbone, were detected, which is in accordance with the literatures.^47^ For PDLLA, a strong carboxyl stretching mode absorption peak around 1750 cm^-1^ was observed, together with peaks around 1450 cm^-1^ and 1380 cm^-1^ ( CH3 asymmetric vibration), and a series of bands between 1000-1300 cm^-1^ ( C-O stretching), in accordance with the literature. ^48^ Moreover, the absence of PVAc signals from zone [i] suggested no inter-layer mixing of PVAc and DAC, thus confirming a clear demarcation between the hydrogel and the primer layer, and the bridging effect of the PVAc primer layer between silicone implant surface and a paintable hydrogel.

### PVAc-mediated coating of silicone implants with sprayable hydrogel (Coseal)

In addition to pre-formed hydrogels like DAC, in situ formed hydrogels that can be applied to substrate surfaces via spraying are also commercially available. Coseal surgical sealant is one of such hydrogels, and its gelation relies on the rapid chemical reactions between 4-arm poly(ethylene glycol) polymers.

After confirming the intact PVAc primer layer on zone [i] and [iv] before hydrogel coating (Figure 7A), Coseal was sprayed and solidified quickly on the implant surface (∼10 seconds), thereby forming a thin coating layer (Figure 7B). Upon the first washing procedure, it was observed that the Coseal layer remained more stable than DAC, since it was not washed away on both zone [i] and [ii] (Figure 7C). This indicated higher structural strength of Coseal hydrogel due to a covalent nature of crosslinking. Meanwhile, the hydrogel applied directly to the silicone surface (zone [ii]) appeared to swell and detach slightly from the surface. A similar yet less significant phenomenon was visible on the PVAc-primed surface (zone [i]). However, after a second washing process, the Coseal hydrogel layer in zone [ii] completely detached from the surface (Figure 7D). On the contrary, the hydrogel layer in zone [i] remained on the PVAc-primed surface despite the formation of wrinkles.

**Figure 7:**
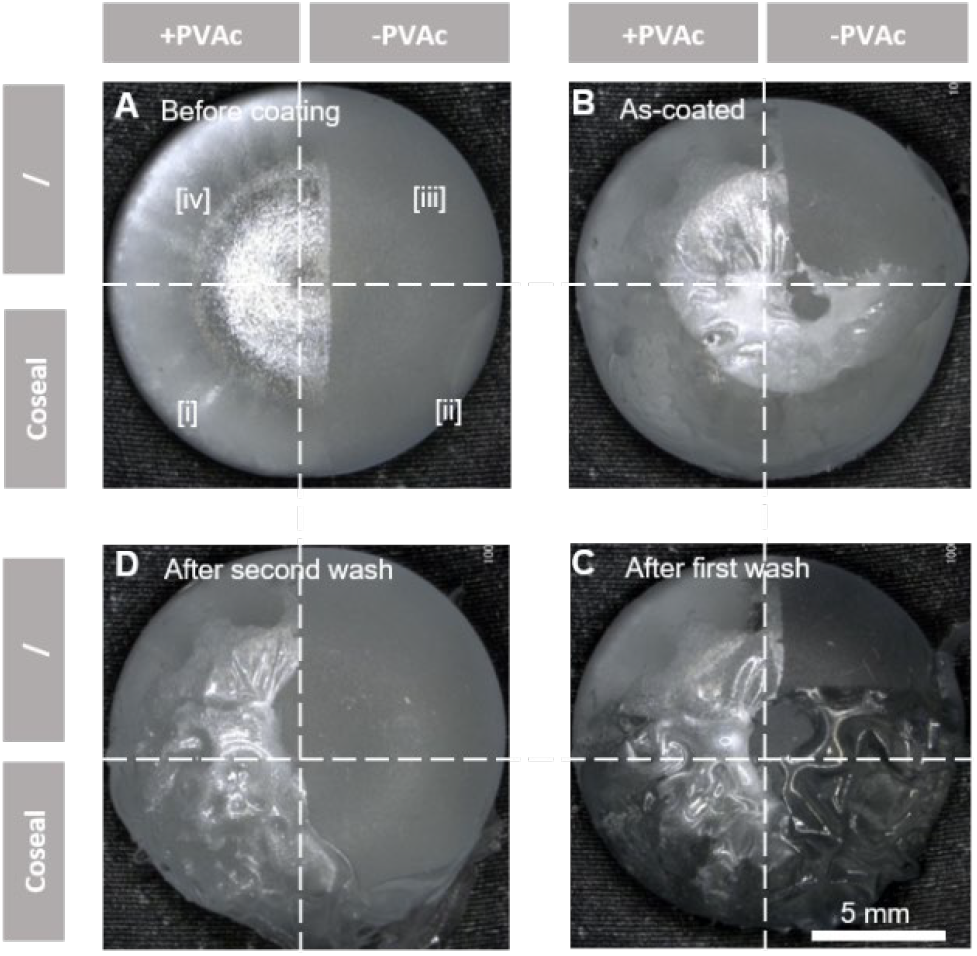
PVAc primer layer mediated coating of silicone implants with sprayable Coseal hydrogel. Silicone implants were divided into four different zones, visualized by dotted white lines. The images were taken before coating with Coseal (A), immediately after *coating (B), after the first (C), and the second wash (D) with PBS*.

The stable coating of sprayable Coseal hydrogel layer in zone [i] was further confirmed by ATR-FTIR analysis (Figure 8). Typical PEG characteristics with absorption bands in the 1100 cm^-1^ region (for ether linkages C-O-C) was observed, accompanied by the signals around 2850 cm^-1^ (C-H stretching vibrations), and a broad absorption band around 3400 cm^-1^ (O-H stretching vibrations from water absorbed in the hydrogel or hydroxyl groups from PEG).^49^ It is noteworthy that the presence of absorption at 1730 cm^-1^ (carbonyl groups, C=O) indicated some exposure of PVAc on the coating surface (Figure 8, zone [i]). This is different from DAC hydrogel coating (Figure 6, zone [i]).

**Figure 8:**
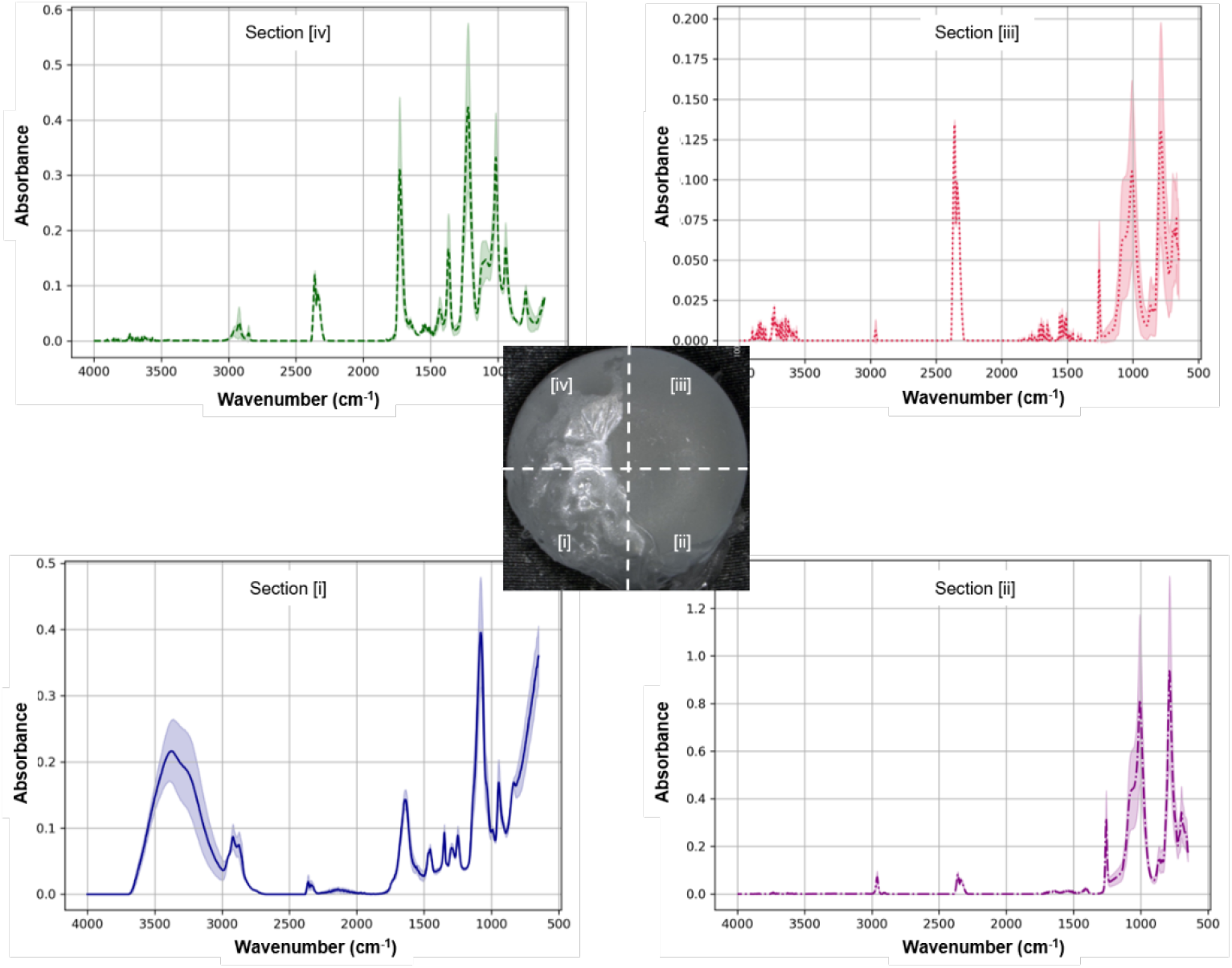
ATR-FTIR spectra of different implant surface zones taken after coating zone [i] and [ii] with Coseal hydrogels (n = 3).

Compared to the pre-formed DAC hydrogel, the precursor solution of Coseal was sprayed onto the implant surface. The liquid nature of this formulation prior to in situ gelation may facilitate dissolution and diffusion of PVAc polymers, thus resulting in the presence of PVAc in the Coseal layer. Conversely, the non-primed surface (zone [ii]) showed the same peaks as the silicone reference surface (zone [iii]). Changes in signal amplitudes occurred between zone [ii] and [iii], which could be attributed to the residues of the Coseal hydrogels in zone [ii]. Unlike the application of DAC via painting, the application of Coseal via spraying appeared to be more challenging, with lower spatial accuracy of the target coating. This resulted in coating more than 50% of the implant surface.

## Conclusion

Hydrogel coating of implants is an attractive technique to avoid intraoperative contamination during implantation, and thus later development of implant-associated infection. The hydrophobic characteristics of silicone surfaces hinder homogenous adhesion of hydrogels on silicone breast implants. To enable stable hydrogel coating, we established a method to use a PVAc primer layer that straightforwardly improves the wettability of silicone surface. This surface priming strategy was proven effective for two commercially available types of hydrogels authorized for internal application, namely, paintable (represented by DAC) and sprayable (represented by Coseal) hydrogels. This method provides opportunities to investigate hydrogel coating of silicone implants, particularly for preventing infection in breast augmentation and reconstruction with implants. Moreover, in a broader sense, the easy application of hydrogels can render implantable biomaterials with a tissue-mimicking interfaces, which could reduce foreign body response.^20^

## Conflicts of interest

The authors declare that there are no conflicts of interest regarding the publication of this paper. All authors have reviewed and approved the manuscript and have agreed to its submission. The authors have no affiliations, financial interests, or personal relationships that could have appeared to influence the work reported in this paper.

## Data availability

The data supporting this article have been included as part of the Supplementary Information.

## Notes

### Competing Interest Statement

The authors have declared no competing interest.

