## supporting information for "Enabling Hydrogel Coating on Silicone Breast Implants with Poly(Vinyl Acetate) Primer Layer"

### Direct hydrogel coating on implant without pre-treatement

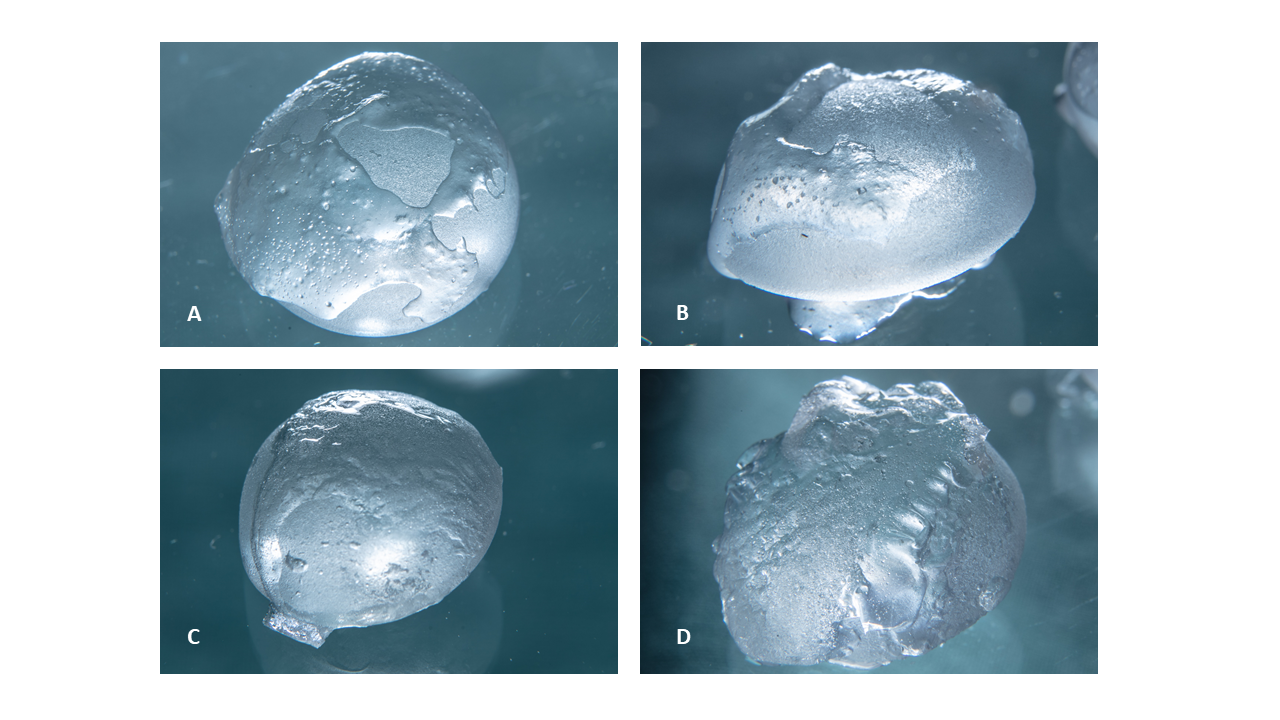

*Figure S1: Silicone implants coated directly with hydrogel. Image A and B: The same silicone implant coated with DAC hydrogel, before (A) and after addition of two drops of NaCl 0,9% (B). Image C and D: Another silicone implant, coated with Coseal hydrogel, before (C) and after addition two drops of NaCl 0,9% (D). Hydration was performed to simulate physiologic surroundings.*

### Step-by-step PVAc priming and surface structure changes

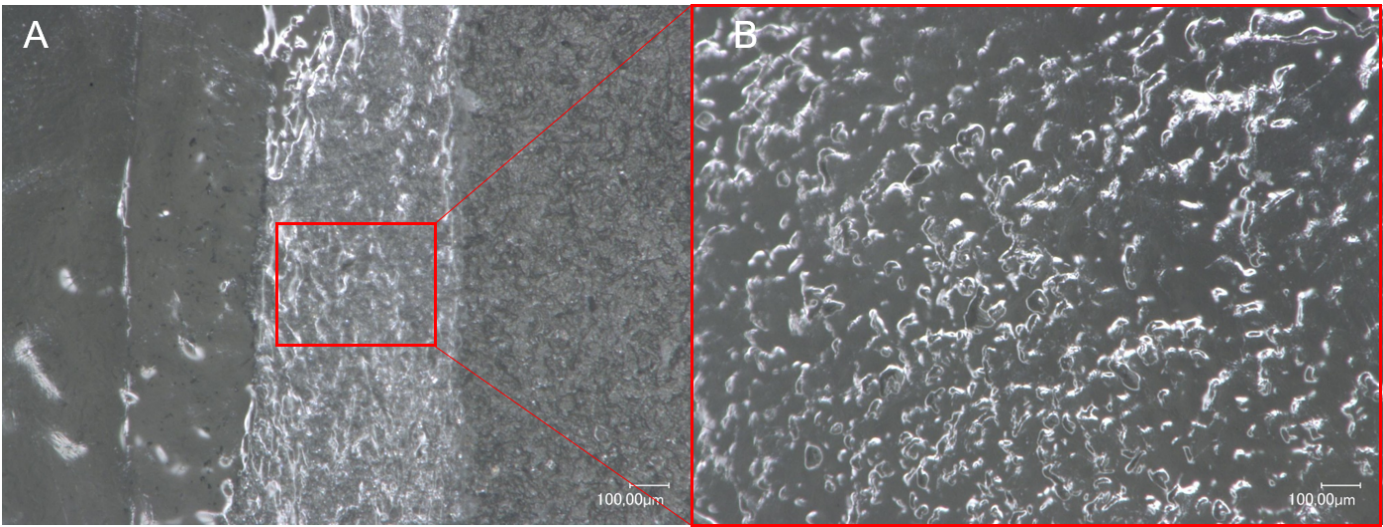

*Figure S2: PVAc primer layer coated on silicone implant. Image A: Section of silicone implant coated with three (left), one (middle), no (right) PVAc priming layer. Image B: Close-up of the one-time coated section. Coating the implant only once leads to insufficient coating and leads to exposure of rough silicone surface structure.*
